# Metacognition, cortical thickness, and tauopathy in aging

**DOI:** 10.1101/2021.10.27.466146

**Authors:** Kailin Zhuang, Xi Chen, Kaitlin E. Cassady, Suzanne L. Baker, William J. Jagust

## Abstract

We investigated two aspects of metacognition and their relationship with cortical thickness and Alzheimer’s Disease (AD) biomarkers, amyloid and tau, in cognitively healthy older adults (N=151). The two metacognition measures were self-appraisal rating of task performance and the difference between self-appraisal rating and actual task performance (appraisal discrepancy). All participants underwent neuropsychological testing and 1.5T structural MRI. A subset (N=66) received amyloid-PET with [11C] PiB and tau-PET with [18F] Flortaucipir. We found that worse performers had lower self-appraisal ratings, but still overestimated their performance, consistent with the Dunning-Kruger effect. Self-appraisal rating and appraisal discrepancy revealed distinct relationships with cortical thickness and AD pathology. Greater appraisal discrepancy, indicating overestimation, was related to thinning of inferior-lateral temporal, fusiform, and rostral anterior cingulate cortices. Lower self-appraisal was associated with higher entorhinal and inferior temporal tau. These results suggest that overestimation could implicate structural atrophy beyond AD pathology, while lower self-appraisal could indicate early behavioral alteration due to AD pathology, supporting the notion of subjective cognitive decline prior to objective deficits.

## 1. Introduction

Metacognition is the awareness and monitoring of one’s own cognitive ability or thought processes (Flavell, 1976; Brown, 1978; Cosentino and Stern, 2005). It is often impaired in patients with Alzheimer’s Disease (AD) (Starkstein, 2014; Rosen et al., 2014; Hallam et al., 2020). Two pathologic features of AD are β-amyloid (Aβ) accumulation and hyperphosphorylated tau aggregation. Both pathologies typically occur decades before the onset of symptoms (Jack et al., 2013). In addition, cortical atrophy in regions affected by AD may be identified almost a decade before reaching the symptomatic stage (Dickerson et al., 2011). Understanding changes in metacognition related to AD pathology during the asymptomatic period might provide clinicians and scientists an important window into the very early behavioral alterations related to AD pathology, improving ability of early detection for AD.

A closely related concept to metacognition within the context of AD research is subjective cognitive decline (SCD), or self-perceived concerns of reduced cognition or memory without objective cognitive impairment (Jessen et al., 2014). SCD is common in the aging population (van Harten et al., 2018) and is proposed as a feature of the earliest detectable behavioral alteration in the AD continuum according to the National Institute of Aging-Alzheimer’s Association research framework (Jack et al., 2018). SCD is associated with altered brain structure (Fan et al., 2018) and function (Chen et al., 2021) as well as Aβ (Perrotin et al., 2017) and tau (Swinford et al., 2018) deposition and hypometabolism (Mosconi et al., 2008; for review, see Wang et al., 2020). However, not all clinically normal elderly whose memory and cognitive functions are actually declining report SCD, and not all individuals with SCD show signs of AD. Traditional neuropsychological methods are not very sensitive to detecting early behavioral deficits in preclinical AD due to individual variability in performance (Saxton et al., 2004), and quantitative measures of metacognition have not been widely applied to investigations of SCD (Molinuevo et al., 2017). Therefore, quantitively examining metacognitive ability and accuracy might give us a new way of identifying and understanding SCD in cognitively healthy individuals.

The goals of the present study were to examine (i) the behavioral characteristics of metacognition, (ii) the structural correlates of metacognition, and (iii) the relationship between AD biomarkers of Aβ and tau and metacognition in a sample of cognitively healthy older adults. We incorporated two measures of metacognition: self-appraisal rating and appraisal discrepancy, which may provide different insights into one’s metacognitive status. As participants completed a series of cognitive tasks, we recorded participants’ self-appraisal rating of task performance in real time as an “online” monitoring measure. We then calculated appraisal discrepancy by subtracting objective performance from self-appraisal as an accuracy measure. We investigated these two metacognitive measures and their relationships with cortical thickness and AD biomarkers. We aimed to explore how disease pathology might be manifested in the subjective assessment of an individual’s ability at a time when objective impairment was not apparent.

## 2. Material and Methods

### 2.1 Participants

A total of 151 cognitively healthy older adults from the Berkeley Aging Cohort Study (BACS) were included in the study. The BACS is an ongoing longitudinal investigation of cognitive and brain changes in cognitively healthy individuals. All participants included in this study were over age 65, had Mini Mental State Examination (MMSE) score >= 25, had neuropsychological testing data, and received 1.5T structural magnetic resonance (MRI) imaging. Sixty six of the 151 participants later received positron emission tomography (PET) imaging for Aβ with ^11^C-Pittsburgh compound B (PiB) and tau with ^18^F-Flortaucipir (FTP).The study was approved by the Institutional Review Board at Lawrence Berkeley National Laboratory (LBNL) and the University of California, Berkeley. Written, informed consent was obtained from all study participants.

### 2.2 Neuropsychological assessment

All participants underwent a standard neuropsychological testing session that assessed episodic memory, working memory, visuospatial ability, executive function, and language. Subclinical depressive symptoms were assessed with the Geriatric Depression Scale (GDS) (Yesavage et al., 1983). For analyses using MRI data to examine cortical thickness and metacognition, the neuropsychological testing session was within six months of the MRI scan. For analyses using PET data to examine AD biomarkers and metacognition, the neuropsychological testing session was within six months of the tau PET scan.

For the present study, we included six cognitive measures that had corresponding online self-appraisal ratings: long delay free recall of the California Verbal Learning Test (Delis et al., 2000), delayed recall of Visual Reproduction (Wechsler, 1997), Digit Span (Wechsler, 1997), number correct in 60 sec of the Stroop test (Golden, 1978), Verbal Fluency (Spreen and Benton, 1977), and Category Fluency. For each task, raw scores were converted to z-scores and then percentile frequencies on a normal distribution. An average objective performance score (in percentile) was calculated by taking the mean of all six task performance percentiles for each subject.

Two metacognitive variables were included in the study: self-appraisal rating and appraisal discrepancy. We used an online performance monitoring paradigm (Perrotin et al., 2012) to assess participants’ metacognitive ability and accuracy. A post-diction self-appraisal rating was recorded after each of the above six tasks. The participant was asked to estimate their task performance on a percentile scale relative to their peers of the same age, sex, and education. An average self-appraisal rating (in percentile) was computed for each participant by taking the mean of all six task self-appraisal ratings. A higher self-appraisal rating indicates better self-perceived performance.

An appraisal discrepancy score was calculated for each task by subtracting the task objective performance score from the participant’s self-appraisal rating. An average appraisal discrepancy score was calculated by taking the mean of all appraisal discrepancy scores of the six tasks. A positive appraisal discrepancy score indicates an overestimation of performance; a negative score indicates underestimation of performance. A score of zero represents perfect accuracy.

### 2.3 MRI acquisition and preprocessing

T1-weighted magnetization-prepared rapid gradient acquisition with gradient echo (MPRAGE) structural MRI scans were acquired for all participants with a 1.5T Siemens Magnetom Avanto scanner at LBNL (voxel size = 1 mm isotropic, repetition time (TR) = 2110 ms, echo time (TE) = 3.58 ms, flip angle (FA) = 15°). All T1 MPRAGE scans were processed using FreeSurfer version 5.3 (https://surfer.nmr.mgh.harvard.edu). Separate MRIs were obtained for the measurement of cortical thickness, and for parcellating the brain for PET data analysis.

### 2.4 PET acquisition and preprocessing

Details of PiB and FTP-PET acquisition were published previously (Ossenkoppele et al., 2016; Schöll et al., 2016). PiB and FTP were synthesized at the Biomedical Isotope Facility at LBNL and all PET scans were acquired on a Biograph 6 Truepoint PET/CT scanner in 3D acquisition mode. Prior to PET scanning, a computed tomography (CT) scan was collected for attenuation correction. Participants were injected with 15 mCi of PiB. 90 minutes of dynamic emission data were acquired and then binned into 35 frames (4 × 15 seconds, 8 × 30 seconds, 9 × 60 seconds, 2 × 180 seconds, 10 × 300 seconds, and 2 × 600 seconds). For tau PET, participants were injected with 10 mCi of FTP and data acquired from 80 to 100 minutes post-injection were binned as 4 × 5 minute frames. PET images were reconstructed using an ordered subset expectation maximization algorithm with weighted attenuation and scatter correction and smoothed with a 4 mm Gaussian kernel. PIB and FTP PET were usually performed on the same day.

For PiB-PET data processing, distribution volume ratio (DVR) values were generated with Logan graphical analysis by calculating the slope from 35 to 90 minutes post-injection and normalized using the cerebellar gray matter as reference region (Logan et al., 1996; Price et al., 2005). A global PiB index was calculated using multiple FreeSurfer derived ROIs as previously described (Mormino et al., 2012) and a global PiB DVR threshold of 1.065 was used to determine PiB positivity (Villeneuve et al., 2015).

For FTP-PET data processing, standardized uptake value ratio (SUVR) images were created based on the mean tracer retention from 80 to 100 minutes post-injection and normalized by the mean tracer retention in the inferior cerebellar gray matter. SUVR images were partial volume corrected (PVC) using the Rousset Geometric Transfer Matrix approach as previously described (Rousset et al., 1998; Baker et al., 2017). We focused on two FreeSurfer parcellated ROIs, entorhinal cortex and inferior temporal cortex, and took the average of the regional FTP SUVR of the left and right hemispheres to create the mean regional FTP SUVR used in the analyses.

### 2.5 Statistical analyses

Statistical analyses were conducted using R (https://www.R-project.org) for cognitive and regional PET data, and FreeSurfer was used for analysis of whole-brain MRI data. First, associations between objective performance and metacognitive measures were explored using Pearson’s correlation. Then, for both metacognitive measures as well as objective cognitive performance, three whole brain vertex-wise analyses were conducted to examine their relationship to cortical thickness, using general linear models in FreeSurfer and adjusting for age, sex, education, and GDS. Multiple comparison correction was applied using Monte Carlo Simulation, with both a liberal *p* value set at *p* < .01 and a stricter *p* value set at *p* < .001. Finally, multiple regressions were used to investigate the relationship between AD biomarker pathology and metacognitive measures as well as objective cognitive performance. All models were adjusted for age, sex, education, and GDS, and all predictors were mean centered to minimize multicollinearity; in these models PiB × FTP interactions were also included because of the well-known acceleration of Aβ on tau effects (Hanseeuw et al., 2019). For models with significant PiB × FTP interactions, we used the Johnson-Neyman procedure to identify the range of global PiB DVR values where regional FTP SUVR had a significant effect on the metacognition measures (Johnson and Fay, 1950; Aiken et al., 1991).

## 3. Results

### 3.1 Demographics

Participants’ demographic characteristics are presented in Table 1. A total of 151 participants (89 female, 62 male) were included in the study, with a mean age of 76.09 years (SD 5.60) and an average of 16.79 years of education (SD 1.92). In the subset of 66 participants who received PET scans, 34 were PiB- and 32 were PiB+, reflecting our efforts to retain PIB+ participants in the study. The PiB+ group was significantly younger than the PiB-group (*p* = .029) but there were no significant group differences in sex, years of education, GDS, MMSE, and neuropsychological tasks performance. There were no significant differences in any variables between the MRI only group and the PET group (all *p* > .1)

**Table 1:** Demographics

|  | MRI Sample<br>N = 151<br>(Mean (SD)) | PET Sample |  |  | Effect size | *p* value<br>(PiB+ vs PiB-) |
| --- | --- | --- | --- | --- | --- | --- |
|  |  | Total<br>N = 66<br>(Mean (SD)) | PiB Negative<br>N = 34<br>(Mean (SD)) | PiB Positive<br>N = 32<br>(Mean (SD)) |  |  |
| Sex = Male (%) | 62 (41.1) | 29 (43.9) | 13 (38.2) | 16 (50.0) | V = 0.09 | 0.475 |
| Age (years) | 76.09 (5.60) | 78.36 (5.29) | 79.74 (6.06) | 76.91 (3.90) | *d* = 0.55 | 0.029 |
| APOE ε4+ (n (%)) | 43/149 (29%) | 20/66 (30%) | 3/34 (8.8) | 17/32 (53.1) | V = 0.45 | < 0.001 |
| Education (years) | 16.79 (1.92) | 16.83 (1.78) | 17.15 (1.62) | 16.50 (1.90) | *d* = 0.37 | 0.14 |
| GDS | 3.32 (2.78) | 3.26 (3.15) | 3.65 (3.42) | 2.84 (2.83) | *d* = 0.26 | 0.304 |
| MMSE | 28.69 (1.35) | 28.73 (1.25) | 29.00 (0.89) | 28.44 (1.50) | *d* = 0.46 | 0.067 |
| CVLT LDFR | 10.42 (3.27) | 10.09 (3.60) | 10.56 (3.72) | 9.59 (3.46) | *d* = 0.27 | 0.28 |
| VRII recall | 53.52 (21.84) | 53.21 (20.02) | 52.74 (21.35) | 53.72 (18.84) | *d* = -0.05 | 0.844 |
| Digit span | 16.11 (3.81) | 15.56 (3.93) | 16.24 (4.40) | 14.84 (3.29) | *d* = 0.36 | 0.152 |
| Stroop correct in 60s | 46.49 (11.43) | 47.88 (10.06) | 48.68 (10.73) | 47.03 (9.39) | *d* = 0.16 | 0.511 |
| Verbal fluency | 46.04 (12.28) | 47.55 (11.27) | 49.38 (11.09) | 45.59 (11.31) | *d* = 0.34 | 0.174 |
| Category fluency | 34.05 (8.40) | 33.09 (8.05) | 33.71 (8.45) | 32.44 (7.69) | *d* = 0.16 | 0.527 |
| COG-MRI interval (days) | 89.84 (44.10) |  |  |  |  |  |
| COG-FTP interval (days) |  | 87.17 (48.35) | 84.88 (48.02) | 89.59 (49.34) | *d* = -0.10 | 0.696 |
| MRI-FTP interval (days)* |  | 1030.94 (1020.37) | 1016.74 (1083.98) | 1046.03 (965.29) | *d* = -0.03 | 0.908 |
| PiB Positive (%) |  | 32 (48.5) |  |  |  |  |

APOE = Apolipoprotein E; GDS = Geriatric Depression Scale; MMSE = Mini-Mental State Examination; CVLT LDFR = California Verbal Learning Test Long Delay Free Recall; VRII = Visual Reproduction Long Delay; FTP = Flortaucipir (\* time between MRI obtained for cortical thickness and FTP PET scan); V = Cramér's V; *d* = Cohen's *d*.

### 3.2 Metacognition and neuropsychological task performance

First, we assessed the relationship between self-appraisal, appraisal discrepancy, and objective neuropsychological task performance. Average task performance had a significant positive association with average self-appraisal ratings (r = .323, *p* < .001) and a significant negative association with average appraisal discrepancy (r = -.621, *p* < .001) (Fig. 1). Thus, individuals with better overall task performance had higher self-appraisal but nevertheless underestimated their performance, whereas those with worse overall task performance had lower self-appraisal but overestimated their performance.

**Fig. 1:**
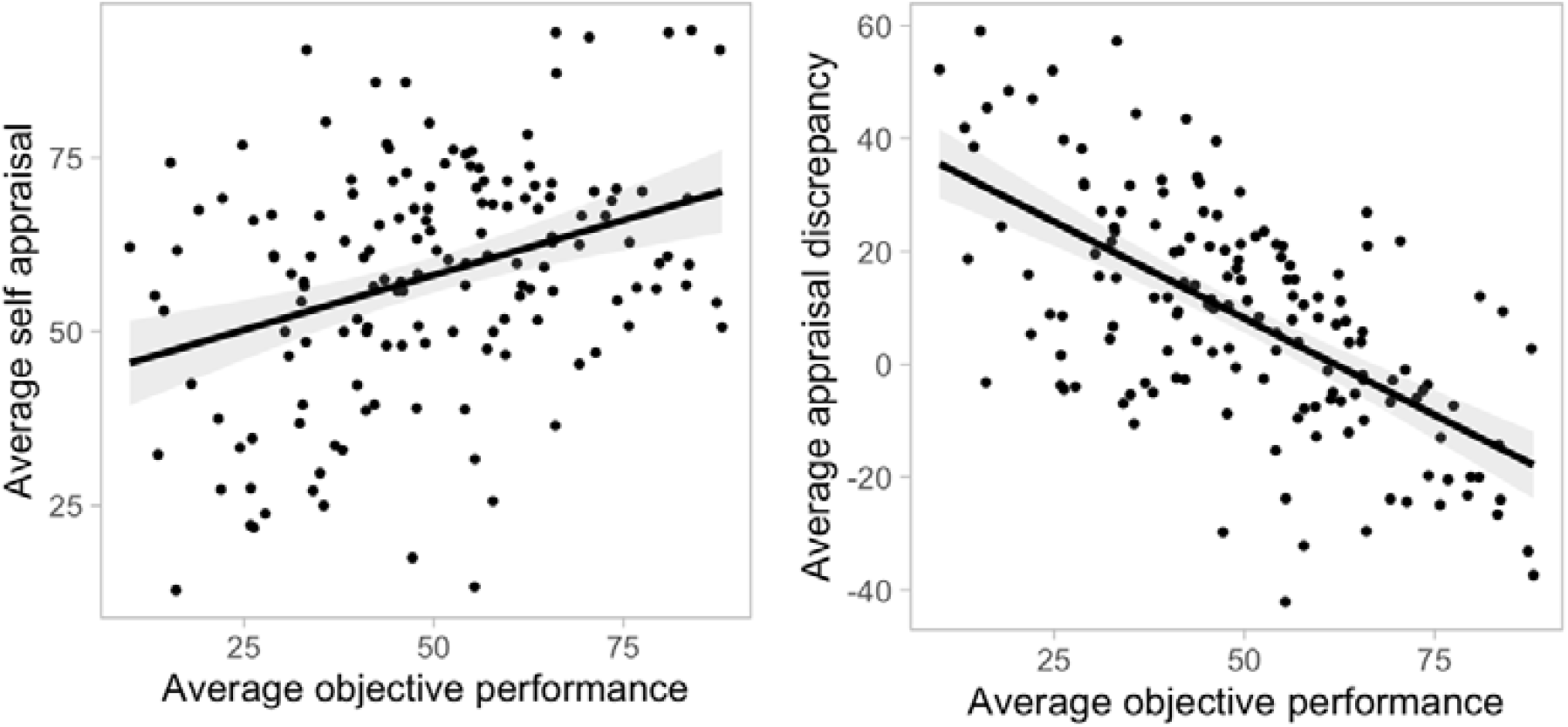
Relationship between metacognitive measures and neuropsychological task performance. Better objective performance was related to higher self-appraisal and lower appraisal discrepancy.

### 3.3 Metacognition and cortical thickness

Next, we explored the relationship between self-appraisal rating, appraisal discrepancy, and cortical thickness, while controlling for age, sex, education, and GDS. The vertex-wise whole-brain cortical thickness analysis revealed a significant negative relationship between average appraisal discrepancy and cortical thickness (Table 2, Fig. 2A, Supplementary Fig. 1A). More overestimation of performance was associated with thinner cortex predominantly in temporal lobe regions, specifically including bilateral inferior temporal, parahippocampal, fusiform, and right superior temporal, as well as inferior parietal cortices and anterior cingulate (*p* < .01, Monte-Carlo simulation). After applying a stricter threshold of Monte-Carlo simulation at *p* < .001, remaining significant clusters include bilateral parahippocampal, left inferior temporal and inferior parietal, and right superior temporal cortices (Table 2). There was no positive association between appraisal discrepancy and cortical thickness. No association was found between self-appraisal ratings and cortical thickness.

**Table 2.** Results of vertex-wise whole brain cortical thickness analyses examining the relationship between cortical thickness and appraisal discrepancy at a liberal and a conservative threshold of Monte Carlo simulation correction for multiple comparisons.

| Max | VtxMax | Size(mm<sup>2</sup>) | TalX | TalY | TalZ | CWP | CWPLow | CWPHi | NVtxs | WghtVtx | Region |
| --- | --- | --- | --- | --- | --- | --- | --- | --- | --- | --- | --- |
| Liberal threshold, Monte Carlo simulation at p<0.01 |  |  |  |  |  |  |  |  |  |  |  |
| Left hemisphere |  |  |  |  |  |  |  |  |  |  |  |
| -3.924 | 130706 | 1106.81 | -52.1 | -26.4 | -17.5 | 0.0001 | 0 | 0.0002 | 1626 | -4654.2 | Inferior temporal |
| -3.741 | 53302 | 926.14 | -38.7 | -47.3 | -13.5 | 0.0001 | 0 | 0.0002 | 1491 | -3760.65 | Fusiform |
| -4.918 | 9460 | 715.14 | -39.6 | -60.8 | 23.5 | 0.0014 | 0.0009 | 0.0019 | 1334 | -3681.99 | Inferior parietal |
| -5.641 | 120176 | 665.22 | -47.5 | -47.7 | 18 | 0.0029 | 0.0022 | 0.0036 | 1474 | -4500.41 | Inferior parietal |
| -5.17 | 31775 | 448.68 | -23 | -19.1 | -23.7 | 0.034 | 0.0317 | 0.0363 | 928 | -2887.39 | Parahippocampal |
| Right hemisphere |  |  |  |  |  |  |  |  |  |  |  |
| -3.561 | 70560 | 1570.56 | 52.8 | -27.8 | -22.1 | 0.0001 | 0 | 0.0002 | 2448 | -5992.8 | Inferior temporal |
| -5.039 | 88831 | 927.27 | 36.3 | -32.7 | 17.5 | 0.0001 | 0 | 0.0002 | 2099 | -5713.16 | Superior temporal |
| -3.421 | 149270 | 707.79 | 13.1 | 27.7 | 28 | 0.0021 | 0.0015 | 0.0027 | 1494 | -3651.81 | Anterior cingulate |
| -4.799 | 127730 | 451.04 | 24.5 | -18.2 | -23.9 | 0.0361 | 0.0337 | 0.0385 | 847 | -2650.38 | Parahippocampal |
| -2.729 | 53317 | 444.37 | 28.2 | -51.5 | -10.7 | 0.0394 | 0.0369 | 0.0419 | 859 | -1937 | Fusiform |
| Conservative threshold, Monte Carlo simulation at p<0.001 |  |  |  |  |  |  |  |  |  |  |  |
| Left hemisphere |  |  |  |  |  |  |  |  |  |  |  |
| -3.924 | 130706 | 480.62 | -52.1 | -26.4 | -17.5 | 0.0001 | 0 | 0.0002 | 707 | -2359 | Inferior temporal |
| -5.17 | 31775 | 249.43 | -23 | -19.1 | -23.7 | 0.0077 | 0.0066 | 0.0088 | 513 | -1851.03 | Parahippocampal |
| -5.641 | 120176 | 214.52 | -47.5 | -47.7 | 18 | 0.0156 | 0.014 | 0.0172 | 447 | -1862.46 | Inferior parietal |
| Right hemisphere |  |  |  |  |  |  |  |  |  |  |  |
| -4.799 | 127730 | 252.7 | 24.5 | -18.2 | -23.9 | 0.0077 | 0.0066 | 0.0088 | 474 | -1725.98 | Parahippocampal |
| -5.039 | 88831 | 175.43 | 36.3 | -32.7 | 17.5 | 0.0361 | 0.0337 | 0.0385 | 461 | -1661.23 | Superior temporal |

Max = maximum $-\log_{10}(p \text{ value})$ in the cluster; VtxMax = vertex number at the maximum; Size = surface area (mm<sup>2</sup>) of cluster; Tal(XYZ) = the talairach coordinate of the maximum; CWP = cluster-wise $p$ value; CWPLow & CWPHi = 90% confidence interval for CWP; NVtxs = number of vertices in cluster; WghtVtx = weight of cluster (size x intensity)

**Fig. 2:**
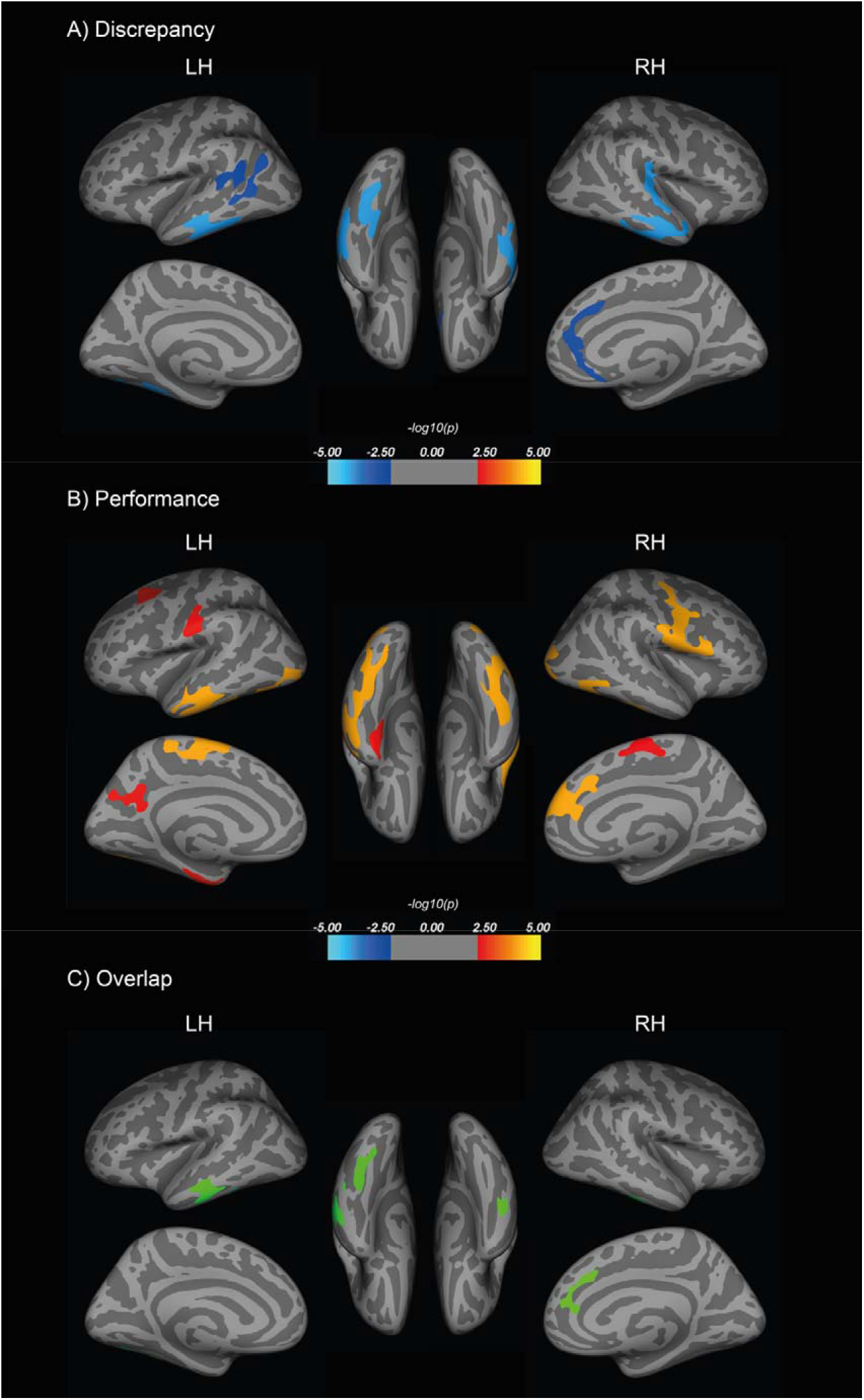
Vertex-wise whole brain analyses were conducted to examine the relationship between cortical thickness and (A) average appraisal discrepancy (overestimation correlated with thinner cortices) and (B) average objective performance, after adjusting for age, sex, education, and GDS (worse objective performance correlated with thinner cortices). The maps shown here used a liberal threshold of cluster-wise Monte Carlo simulation correction at *p* < 0.01. Overlapping regions are presented in (C).

We also found a significant relationship between better objective cognitive performance and greater thickness in a wide range of cortices, including bilateral precuneus, left middle temporal, middle frontal, fusiform, superior parietal, paracentral and postcentral, and right inferior temporal, superior frontal, lateral occipital, paracentral, and precentral regions (Fig. 2B, Supplementary Fig. 1B). Next, we applied the discrepancy cortical thickness map onto the objective cognitive performance cortical thickness map to generate an overlap of identified cortical regions in common, including bilateral inferior temporal and fusiform, left middle temporal, and right superior frontal and rostral anterior cingulate cortices (Fig. 2C). Cortical thinning in these areas was related to worse objective performance and more overestimation of performance.

### 3.4 Metacognition, global Aβ, and regional tau

We investigated the relationship between self-appraisal rating, global PiB DVR, and entorhinal or inferior temporal FTP SUVR, while controlling for age, sex, education, and GDS (Table 3, models 1 & 2). Higher entorhinal FTP SUVR was associated with lower self-appraisal (*p* = .018). There was a significant interaction between global PiB DVR and entorhinal FTP SUVR (*p* = .019) (model 1, Fig. 3A, B), such that higher global Aβ level increased the effect of higher entorhinal tau on lower average self-appraisal ratings. The effect of entorhinal FTP SUVR on self-appraisal became significant after global PiB DVR reached a value of 1.11 (corresponding to 16 Centiloids). Higher GDS (*p* < .001) and lower global PiB DVR (*p* = .003) were also significantly associated with lower self-appraisal ratings. There was also a significant negative association between average self-appraisal rating and inferior temporal FTP SUVR (*p* = .045; model 2, Fig. 3C). In this model, higher GDS (*p* < .001) and lower PiB DVR (*p =* .037) were significantly related to lower self-appraisal, but there was no significant interaction between global PiB DVR and inferior temporal FTP SUVR on average self-appraisal rating.

**Table 3:** Regression statistics estimating appraisal discrepancy and self-appraisal from EC and IFT FTP and covariates

| Covariates | Estimate | Std. Error | t value | p value | sig |
| --- | --- | --- | --- | --- | --- |
| *Model 1. Self-appraisal and EC FTP* |  |  |  |  |  |
| (Intercept) | 0.537 | 2.225 | 0.241 | 0.810 |  |
| EC FTP | -20.138 | 8.265 | -2.437 | 0.018 | * |
| Age | 0.653 | 0.335 | 1.950 | 0.056 | † |
| sexMale | 1.943 | 3.353 | 0.580 | 0.564 |  |
| GDS | -1.990 | 0.513 | -3.882 | 0.000 | *** |
| Education | -1.300 | 0.918 | -1.415 | 0.162 |  |
| PiB DVR | 31.915 | 10.462 | 3.051 | 0.003 | ** |
| PiB x FTP | -64.738 | 26.714 | -2.423 | 0.019 | * |
| *Model 2. Self-appraisal and IFT FTP* |  |  |  |  |  |
| (Intercept) | 0.003 | 2.397 | 0.001 | 0.999 |  |
| IFT FTP | -24.421 | 11.939 | -2.046 | 0.045 | * |
| Age | 0.548 | 0.342 | 1.603 | 0.114 |  |
| sexMale | 2.292 | 3.582 | 0.640 | 0.525 |  |
| GDS | -2.044 | 0.539 | -3.792 | 0.000 | *** |
| Education | -1.711 | 0.978 | -1.749 | 0.086 | † |
| PiB DVR | 22.487 | 10.554 | 2.131 | 0.037 | * |
| PiB x FTP | -74.157 | 45.029 | -1.647 | 0.105 |  |
| *Model 3. Appraisal discrepancy and EC FTP* |  |  |  |  |  |
| (Intercept) | -4.912 | 2.971 | -1.654 | 0.104 |  |
| EC FTP | 15.102 | 11.035 | 1.369 | 0.176 |  |
| Age | 0.464 | 0.447 | 1.039 | 0.303 |  |
| sexMale | 15.422 | 4.477 | 3.445 | 0.001 | ** |
| GDS | -1.934 | 0.685 | -2.825 | 0.006 | ** |
| Education | -3.497 | 1.226 | -2.852 | 0.006 | ** |
| PiB DVR | 21.578 | 13.969 | 1.545 | 0.128 |  |
| PiB x FTP | -86.759 | 35.667 | -2.432 | 0.018 | * |
| *Model 4. Appraisal discrepancy and IFT FTP* |  |  |  |  |  |
| (Intercept) | -6.070 | 3.159 | -1.922 | 0.060 | † |
| IFT FTP | -1.992 | 15.733 | -0.127 | 0.900 |  |
| Age | 0.684 | 0.451 | 1.517 | 0.135 |  |
| sexMale | 15.555 | 4.721 | 3.295 | 0.002 | ** |
| GDS | -2.097 | 0.710 | -2.953 | 0.005 | ** |
| Education | -3.364 | 1.289 | -2.611 | 0.011 | * |
| PiB DVR | 17.878 | 13.908 | 1.285 | 0.204 |  |
| PiB x FTP | -56.129 | 59.339 | -0.946 | 0.348 |  |

\*\*\* $p < 0.001$ , \*\* $p < 0.01$ , \* $p < 0.05$ , † $p < 0.1$ ; EC = Entorhinal Cortex; IT = Inferior Temporal; FTP = Flortaucipir; SUVR = Standardized Update Value Ratio; GDS = Geriatrics Depression Scale; PiB = Pittsburgh compound B; DVR = Distribution Volume Ratio

**Fig. 3:**
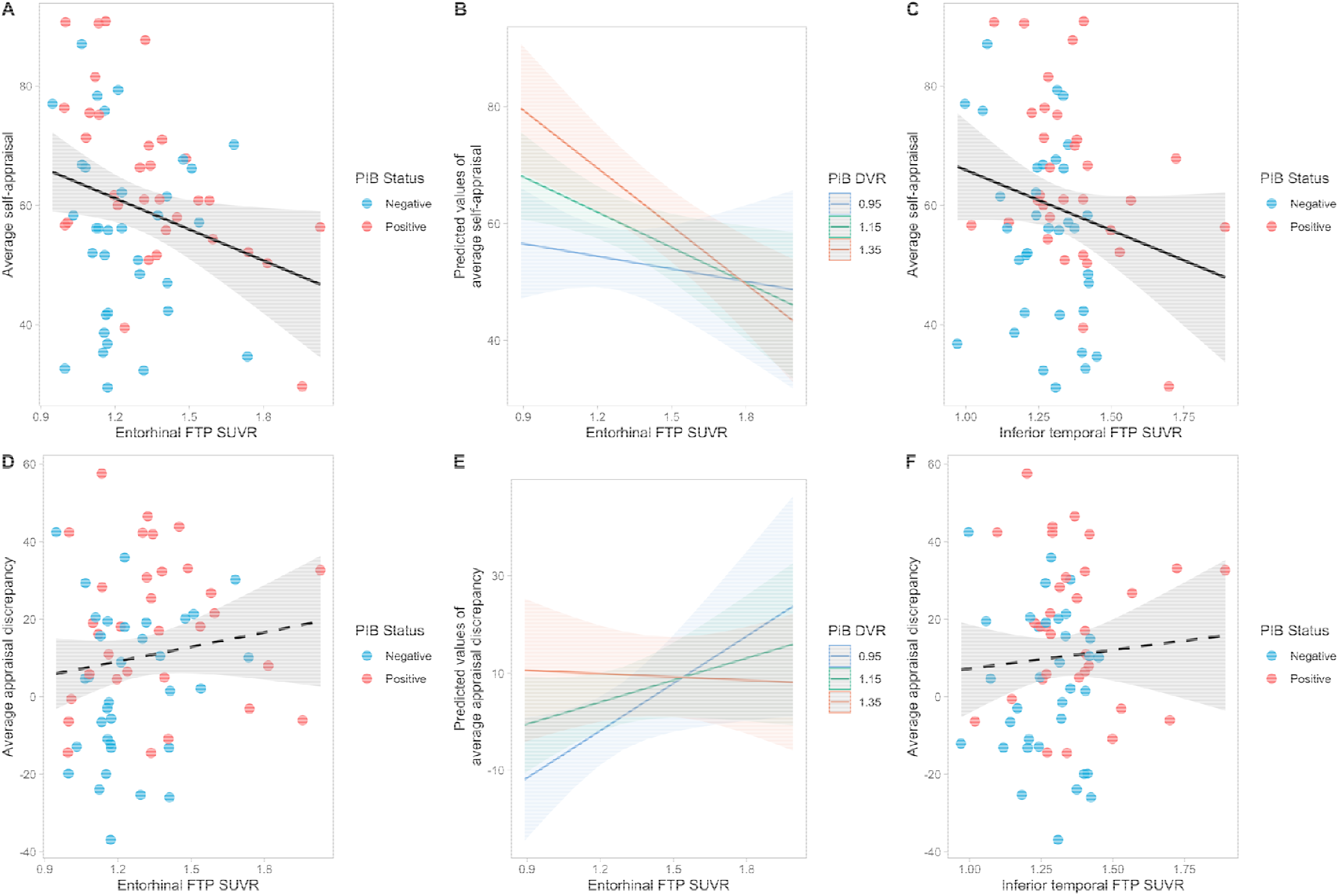
Relationship between metacognitive measures and AD biomarkers, adjusting for age, sex, education, and GDS. Higher entorhinal FTP SUVR was significantly associated with lower average self-appraisal ratings (A), especially in individuals with higher global PiB DVR (B). Higher inferior temporal FTP SUVR was related to lower average self-appraisal ratings (C) but there was no significant interaction between global PiB DVR and inferior temporal FTP SUVR. (D) shows the relationship between entorhinal FTP SUVR and average appraisal discrepancy stratified by PiB positivity. There was a significant interaction between global PiB DVR and entorhinal FTP SUVR: higher entorhinal tau was related to greater average appraisal discrepancy (overestimation) but in individuals with lower amyloid level; this interaction is unlikely to be biologically significant (see text) (E). There was no significant association between inferior temporal FTP SUVR and average appraisal discrepancy (F). FTP = Flortaucipir; SUVR = Standardized Update Value ratio; GDS = Geriatrics Depression Scale; PiB = Pittsburg compound B; PiB positivity DVR threshold = 1.065

Next, we examined the association of appraisal discrepancy, global PiB DVR, and entorhinal/inferior temporal FTP SUVR, while controlling for the same covariates (Table 3, models 3 & 4). There was no significant main effect of entorhinal FTP SUVR on appraisal discrepancy, but a statistically significant interactive effect between global PiB DVR and entorhinal FTP SUVR (*p* = .018) (Fig. 3D, E). We found that the effect of entorhinal FTP SUVR on appraisal discrepancy was predicted to become significant only if global PiB DVR was lower than 1.02 or greater than 1.94. Given the global PiB DVR range 0.98 to 1.89 in our sample, it appears that this interaction was likely to be irrelevant across the majority of PiB values. In addition, being a male (*p* = .002), lower GDS score (*p* = .005), and lower education (*p* = .006) were also found to be significantly associated with a tendency towards overestimation. In the inferior temporal tau model, no significant associations were found between average appraisal discrepancy and inferior temporal FTP SUVR, and there was no interactive effect between global PiB DVR and inferior temporal FTP SUVR on average appraisal discrepancy. However, the significant effects of sex (*p* = .002), GDS (*p* = .005), and education (*p* = .011) remained (model 4, Fig. 3F).

Finally, we examined the relationship between average objective performance and AD biomarkers while adjusting for the same covariates (Supplementary Table 1). Higher entorhinal FTP SUVR was significantly related to lower average objective performance (*p* = .001, Supplementary Fig. 2A) but no significant interaction between global PiB DVR and entorhinal FTP SUVR was found. There was a negative, but statistically insignificant relationship between inferior temporal FTP SUVR and performance (*p* = .140, Supplementary Fig. 2B).

## 4. Discussion

In this study, we investigated the relationships between metacognition and objective performance, brain structure, and AD pathology in a community sample of cognitively healthy older individuals. We used two measures of metacognition: an online self-appraisal rating directly recording participants’ estimation of task performance and an appraisal discrepancy calculated to quantify the extent of over- and underestimation of task performance. We found that worse performers tended to have lower self-appraisal yet still overestimated their cognitive performance, consistent with the notable Dunning-Kruger effect where individuals with worse abilities tended to overestimate their performance and ability (Kruger and Dunning, 1999). The two metacognition measures revealed distinct associations with brain morphology and AD biomarkers. Greater appraisal discrepancy, indicating overestimation of performance, was related to cortical thinning in temporal, anterior cingulate, and parietal cortices. These regions overlapped, to some extent, with brain regions where cortical thinning was related to worse objective performance. However, appraisal discrepancy did not have apparent relationships with regional tau or global Aβ. On the other hand, self-appraisal as a measure of metacognitive awareness was sensitive to AD pathology, but not cortical atrophy. Specifically, lower self-appraisal ratings related to higher inferior temporal tau, as well as higher entorhinal tau in individuals with higher global Aβ deposition. This suggests that even in cognitively healthy older adults, structural differences in temporal, parietal, and medial regions may underlie inaccurate metacognitive appraisal, and that higher AD pathology could be manifested by lower self-appraisal ratings.

Previous research has shown that impaired metacognitive ability is associated with reduced right posterior cingulate and medial prefrontal cortical thickness in a cognitively diverse sample of older adults (Bertrand et al., 2018). Here, we report that in cognitively healthy older individuals, worse objective performance and greater overestimation of performance were related to cortical thinning in inferior-lateral temporal and fusiform and rostral anterior cingulate cortices. Young and older adults with cortical thinning in these areas show reduced memory performance (Busovaca et al., 2016), and may also have worse metacognitive monitoring (Kruger and Dunning, 1999). Decreased cortical thickness of medial and inferior temporal gyri has been shown to be related to increased risk and progression towards AD (Dickerson et al., 2009). Moreover, inferior temporal and fusiform regions form part of the anterior-temporal network, which is particularly vulnerable to early tau deposition (Lowe et al., 2018; Maass et al., 2019), and inferior temporal tauopathy is associated with cognitive impairment (Johnson et al., 2016). We also found greater overestimation to be related to parahippocampal thinning, an area involved in retrospective metamemory judgements (Vaccaro and Fleming, 2018). However, areas where reduced cortical thickness was associated with appraisal discrepancy in our study are not limited to the memory domain. Fusiform morphology is implicated in metacognitive assessment of motor functions. A previous study examining metacognitive ability with a visuomotor task in healthy adults found that higher grey matter volume in the right fusiform gyrus was associated with greater metacognitive sensitivity (Sinanaj et al., 2015). A recent study found that in young adults, greater left fusiform grey matter volume was related to greater overestimation of handwriting quality, another visuomotor function (Li et al., 2021). These findings suggest that while appraisal discrepancy is related to frontal and temporal lobe cortical atrophy, this measure is not specific to AD pathology but rather reflects overall decreased metacognition and functions beyond memory.

The right anterior cingulate has been demonstrated to be involved in error detection (Carter, 1998) and error information processing (Holroyd et al., 2004). The prefrontal and posterior cingulate cortex are involved in self-reflective thought processes (Johnson et al., 2002). Several studies have found the right midline structures to be involved in self-awareness processes (Bertrand et al., 2018; Muñoz-Neira et al., 2019). Moreover, higher levels of anosognosia correlated with greater atrophy and hypometabolism in the left dorsal anterior cingulate in AD patients (Guerrier et al., 2018). Decreased grey matter volume in anterior cingulate and fusiform is related to anosognosia in memory and non-memory domains in MCI and AD patients (Valera-Bermejo et al., 2020). Together, these structural findings suggest that as these brain regions become more atrophic in aging, objective performance deteriorates, self-appraisal ability worsens, and error monitoring capability declines.

In contrast to appraisal discrepancy, we found effects of AD pathology on self-appraisal rating: both higher entorhinal and inferior temporal tau were associated with lower self-appraisal ratings. Consistent with a previous finding (Buckley et al., 2017), decreased self-appraisal of performance was related to increased entorhinal tau burden. We also found an interaction between entorhinal tauopathy and global Aβ level on self-appraisal with higher Aβ exacerbating the tau effect, an effect not observed in Buckley et al (2017). One possible explanation is that even though both studies included cognitively healthy older adults and investigated cognitive deficit awareness, the main variable of interest in our study tapped into the online metacognitive appraisal process, rather than global subjective cognitive concerns, which may be more sensitive to more progressed tau pathology. This is partly supported by the non-significant interactive effect in the inferior tau model: high tau buildup in the inferior temporal lobe is already associated with positive global Aβ deposition (Sanchez et al., 2021) and that Aβ threshold might not play a significant role in this relationship. This suggests that lower self-appraisal, indicating subjective cognitive decline, could be an early indicator of more pronounced AD pathology in the cognitively healthy.

Our findings add to the previous studies investigating AD biomarkers and memory awareness in individuals with MCI (Therriault et al., 2018), anosognosia (Guerrier et al., 2018), and unimpaired individuals (Vogel et al., 2017; Buckley et al., 2017; Chen et al., 2019; Vannini et al., 2019), by quantitively examining two aspects of metacognition and their relations to AD pathology and cortical structure. They appear to reveal a double dissociation where appraisal discrepancy was related to cortical atrophy, but not AD pathology, while self-appraisal was associated with AD pathology, but not cortical atrophy. The intricate relationships between metacognition and pathology suggest a potential timeframe with regard to the differences in the sensitivity of subjective cognitive awareness measures in the detection of cognitive impairment in preclinical AD. Overestimation was related to cortical thinning in temporal, parietal, and anterior cingulate regions, reflecting a metacognitive deficit accompanying lower objective performance. However, the missing relationship with AD biomarkers suggests that this measure is not specific to AD, but likely reflects individual differences and age-related changes in higher order cognitive processes. In contrast, lower self-appraisal ratings were more sensitive to AD pathology. Aβ+ and high tau individuals had lower individual appraisal ratings, which may reflect a more accurate metacognitive process in preclinical AD when behavioral deficits begin to emerge, further suggesting the validity of subjective complaints of worse cognition in early AD. Together, these findings suggest that in the practice of examining subjective cognitive concerns of older adults, using a variety of metacognitive measures might help track SCD status in cognitively healthy individuals and better identify those who may or may not be developing AD pathology.

This study has its limitations. First, while our study highlights the use of online metacognitive measures, having more comprehensive offline questionnaires examining metacognitive abilities would be beneficial to further examine the relationship between metacognition and cognition along with AD biomarkers. Second, the convenience sampling nature of the study resulted in the study demographics being highly educated, affluent, and mostly of white European descent, which does not reflect the demographic profile of the general aging population. In addition, there are few participants in this study sample of cognitively healthy older individuals who fall in the category of having both high amyloid and high tau. Future research could investigate metacognitive abilities and accuracy in different subgroups of cognitively healthy individuals with varying levels of amyloid and tau deposition to clarify and add onto this relationship. Lastly, we are limited in making inferences about the direct relationships between tauopathy and cortical atrophy with metacognition because we do not have tau PET at the same time of the cortical thickness analyses. Cross-sectional snapshots of metacognitive appraisal, performance, and neuropathology are insufficient to understand the intricacy of self-awareness in aging. Future studies should examine longitudinal online metacognitive measures and further explore their relationship to the progression of AD pathology and structural atrophy.

In conclusion, our findings demonstrated that overestimation of performance was associated with cortical thinning in a range of temporal and frontal regions which were also related to poorer objective performance in cognitively healthy older adults. Having higher tau deposition was related to lower self-appraisal in performance which may reflect an increased awareness of performance deficits in individuals with AD pathology.

## Acknowledgements

This work was supported by National Institutes of Health Grants AG034570 and AG062542. Avid Radiopharmaceuticals enabled the use of the 18F-Flortaucipir tracer but did not provide direct funding and were not involved in data analysis or interpretation.

