## Supplemental materials for "Metacognition, cortical thickness, and tauopathy in aging"

**Supplementary material**

Supplementary Table 1 Regression statistics estimating average objective performance from EC and IT FTP and covariates

| Covariates | Estimate | Std. Error | t value | p value | sig |
| --- | --- | --- | --- | --- | --- |
| *Model 1. Performance and EC FTP* | | |  |  |  |
| (Intercept) | 5.449 | 2.748 | 1.983 | 0.052 | † |
| EC FTP | -35.240 | 10.207 | -3.452 | 0.001 | ** |
| Age | 0.189 | 0.414 | 0.456 | 0.650 |  |
| sexMale | -13.478 | 4.141 | -3.255 | 0.002 | ** |
| GDS | -0.056 | 0.633 | -0.089 | 0.929 |  |
| Education | 2.198 | 1.134 | 1.938 | 0.058 | † |
| PiB DVR | 10.337 | 12.921 | 0.800 | 0.427 |  |
| PiB x EC FTP | 22.021 | 32.991 | 0.667 | 0.507 |  |
| *Model 2. Performance and IT FTP* | | |  |  |  |
| (Intercept) | 6.073 | 3.008 | 2.019 | 0.048 | * |
| IT FTP | -22.429 | 14.978 | -1.498 | 0.140 |  |
| Age | -0.135 | 0.429 | -0.315 | 0.754 |  |
| sexMale | -13.263 | 4.494 | -2.951 | 0.005 | ** |
| GDS | 0.053 | 0.676 | 0.079 | 0.937 |  |
| Education | 1.654 | 1.227 | 1.348 | 0.183 |  |
| PiB DVR | 4.609 | 13.241 | 0.348 | 0.729 |  |
| PiB x IT FTP | -18.028 | 56.490 | -0.319 | 0.751 |  |

*** *p* < 0.001, ** *p* < 0.01, * *p* < 0.05, † p < 0.1; EC = Entorhinal Cortex; IT = Inferior Temporal; FTP = Flortaucipir; SUVR = Standardized Update Value Ratio; GDS = Geriatrics Depression Scale; PiB = Pittsburg compound B; DVR = Distribution Volume Ratio

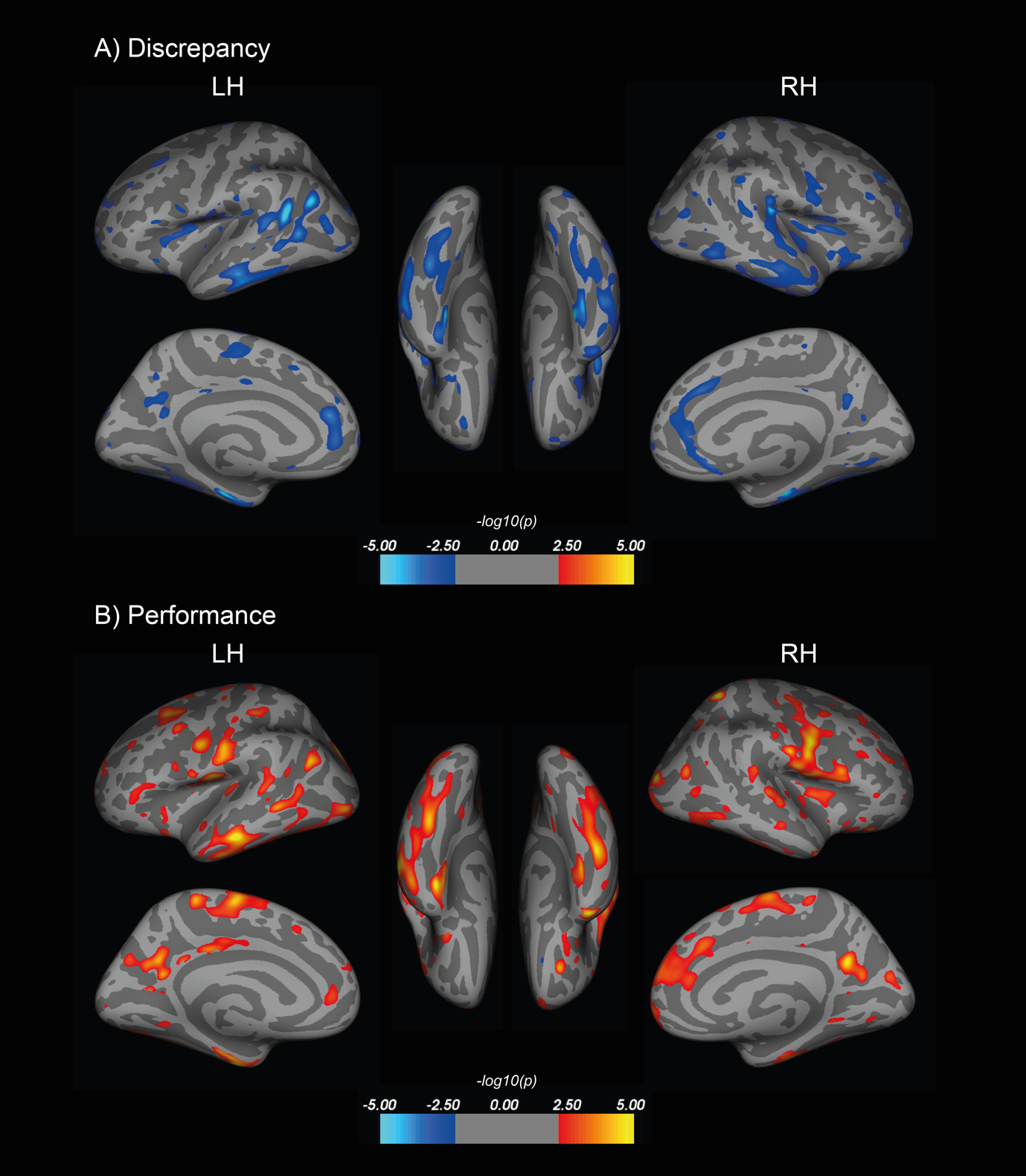

Supplementary Fig. 1: Uncorrected results of vertex-wise whole brain analyses examining the relationship between cortical thickness and (A) average appraisal discrepancy and (B) average objective performance, while adjusting for age, sex, education, and GDS.

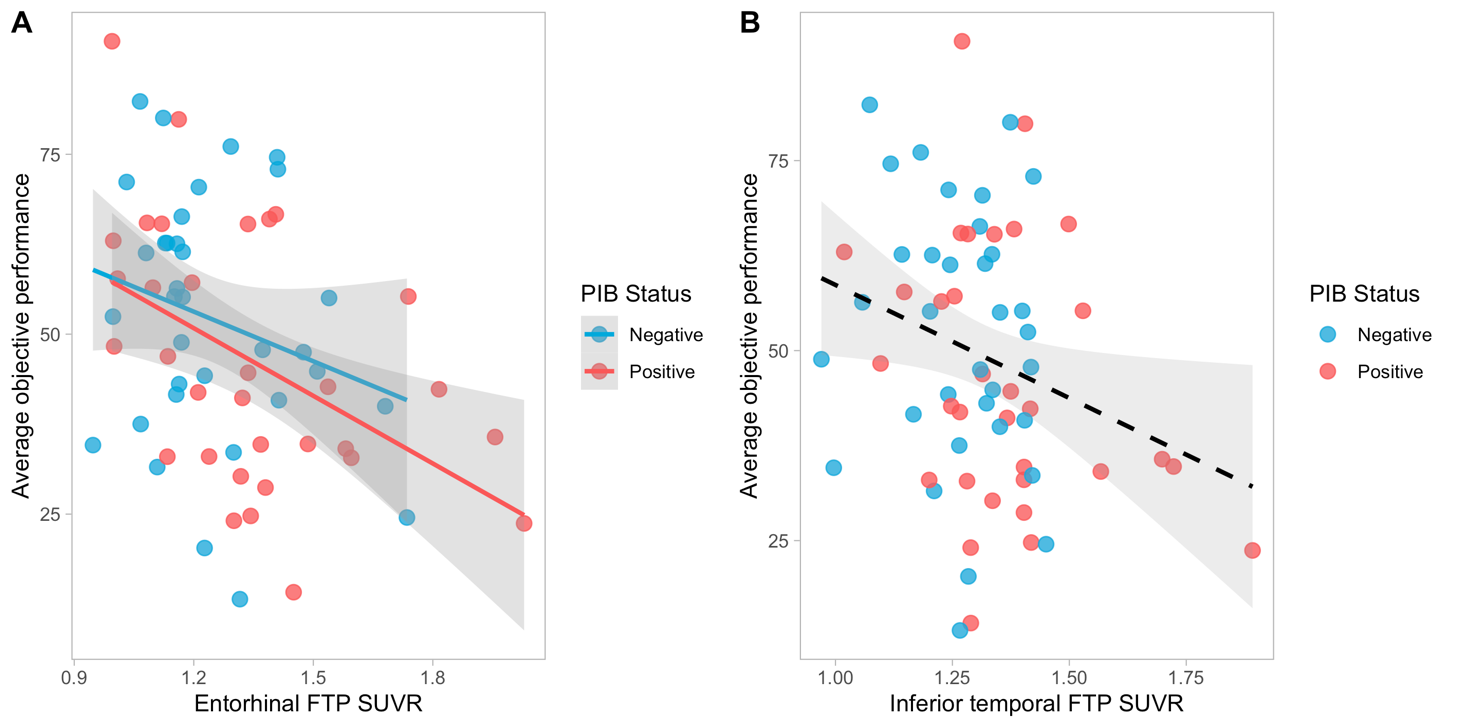

Supplementary Fig. 2: Relationship between average objective performance and AD biomarkers, adjusting for age, sex, education, and GDS. Higher entorhinal FTP SUVR was significantly related to lower average objective performance but no significant interaction between global PiB DVR and entorhinal FTP SUVR was found (A). There was no significant association between inferior temporal FTP SUVR and average objective performance (*p* = .140) (B).

FTP = Flortaucipir; SUVR = Standardized Update Value ratio; GDS = Geriatrics Depression Scale; PiB = Pittsburg compound B; PiB positivity DVR threshold = 1.065
